# Bio-Babel: autonomous cross-language reconstruction of agent-ready computational biology ecosystems

**DOI:** 10.64898/2026.08.30.743167

**Authors:** Nianping Liu, Xuanzhi Chen, Miao Cui, Xiaoying Liao, Xiaoke Song, Sairam Pantham, Weize Xu, Xiaojie Qiu

## Abstract

Scientific software is often locked in its source language, callable from others but not natively extensible or built upon. We present Bio-Babel (biobabel.stanford.edu), an autonomous framework that rebuilds software natively in target ecosystems and makes it agent-callable, shipping each package with an agent-readable contract of its usage. Reconstructing a hierarchical, interdependent R-to-Python single-cell stack, Bio-Babel reproduced the originals faithfully and revealed pancreatic differentiation dynamic defects under graded SWI/SNF loss.

## Main

Scientific software is often locked in the language ecosystem in which it was originally written. R’s grid, the abstract layout machinery beneath the influential grammar-of-graphics (GOG) ggplot2 package^1^ and its many extensions, remains only in R, and foundational tools in single cell genomics such as Monocle 2^2^ have become unmaintained as their original developers and dependencies moved on. Meanwhile, biological analyses, particularly single cell and spatial genomics analyses, are increasingly centered on Python thanks to its better design (computational efficiency, memory management, parallelism and scale), native integration with the machine learning ecosystem, and the active development of the scverse ecosystem^3^ and others. Meanwhile, large-language-model (LLM) agents are increasingly used for data science by retrieving existing packages, composing them and writing new methods on top^4,5^. An agent can still reach across languages through a bridge such as rpy2, but doing so is brittle and can not be recomposed or extended natively. Native full re-implementation of a single package in a new language is often hard, because the behaviour that matters lies not only in a package’s function signatures but also in its object system, data containers, numerical conventions and ecosystem dependencies. Thus, a literal re-implementation can run while silently diverging from the original without joint implementation of supporting dependencies beyond the package itself. Furthermore, without consideration of the target language ecosystem, it will sit outside the idioms of the target language ecosystem it is supposed to belong to.

To close these gaps, here we developed Bio-babel (Fig. 1), a multi-agent framework that reconstructs software natively in another ecosystem and ships it with the operating knowledge an agent needs to use it. To achieve this, two separate requirements must be satisfied: (1) fidelity, that is the rebuilt package should behave like the original version, and (2) usability, that is, an agent that has never seen it should be able to use it correctly. Bio-babel meets both requirements by unifying two interconnected agent-driven systems(Fig. 1a): a build system that reconstructs the software, and an interface system that serves its operating knowledge to the agents.

**Figure 1.**
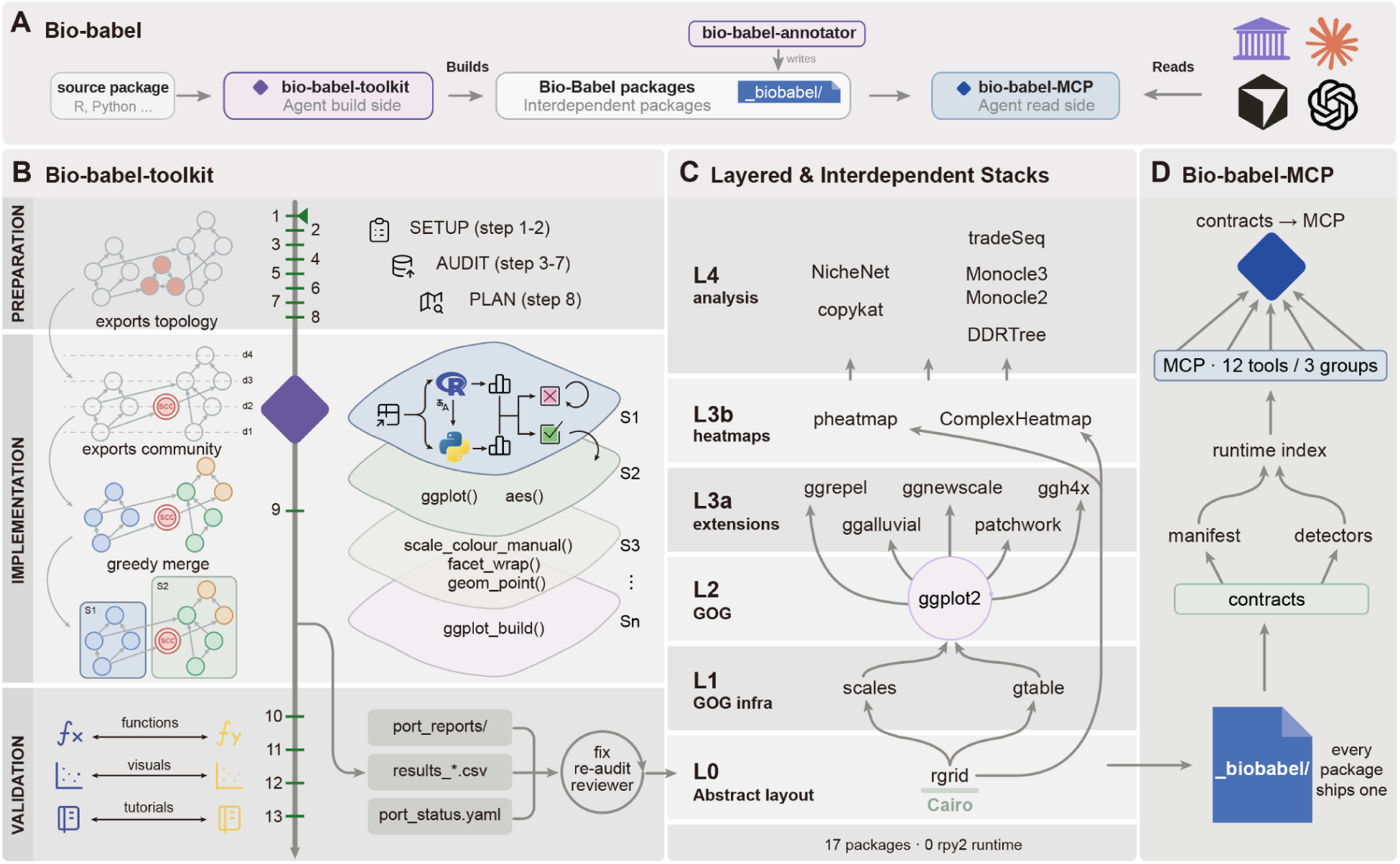
The Bio-Babel translation and execution framework. (**A**) Bio-Babel jointly enables translation of software ecosystems from one language to another and effective agent-calling through Model Context Protocol (MCP) based package registration. (**B**) Schematic of the Bio-Babel-Toolkit workflow, consists of 13 consecutive steps, indicated along the vertical arrow, grouped into preparation (steps 1-8), implementation (step 9) and validation (steps 10-13). The package of interest is first parsed to create a dependency graph between different functions where each node corresponds to a function while the directional edge indicates the dependencies between functions. Circularly connected components in the graph are firstly grouped into strongly connected components (SCC). At step 9 the export-dependency graph is merged through a greedy merging algorithm to form different dependency-ordered translation slices, e.g. S1, S2, etc, each translated through an diagnose-and-test loop, either in tandem (if slices are dependent) or in parallel (if slices are independent), that iterates until equivalence between the translated and original implementation is reached. From steps 10 to 13, the validation system reviews functions, renders visuals and runs tutorials to confirm the function and package translation accuracy. (**C**) The layered workflow to translate a hierarchical R-based ecosystem to Python ecosystem. L0, abstract-layout layer; L1, grammar-of-graphics (GOG) infrastructure layer; L2, the GOG grammar layer; L3a, grammar extensions layer; L3b, grid-direct heatmaps layer; L4, the layer of single-cell analysis methods (for example, monocle 2, tradeSeq). The arrows denote code dependencies. The converted whole stack runs with Python natively without rpy2 at runtime. (**D**) The package registration and contract system to allow convenient calling of the converted packages. Every converted package ships a *_biobabel*/ contract, where two entry-point groups (manifest, detectors) register into a registry serving the agent as a read-only MCP surface.

The build system, Bio-Babel-Toolkit, drives coding agents through a 13-step workflow in three phases: preparation (steps 1-9), implementation (step 9) and validation (steps 10-13) (Fig. 1b and Supplementary Fig. 1a). A controller orchestrates the agentic loop and initializes a local report tree that synchronizes state between the live agents and on-disk progress, so progress lives on disk rather than in the model’s context and survives an agent after losing the state (Supplementary Fig. 1b). Crucially, implementation operates on the package level rather than the function level: it resolves the entire export surface into a dependency graph and partitions it into dependency-ordered slices, each built and tested only after its dependencies. Finally, the port can enter an iterative re-audit step, in which each export is re-checked against its original and any remaining discrepancies are fed back for correction (Supplementary Fig. 1c). On the other hand, Bio-babel-MCP, the interface system, is a decentralized Model Context Protocol (MCP) server that assembles these contracts from each package through Python entry-point groups(Fig. 1d and Supplementary Fig. 2, Methods). This server is a built-in, decentralized broadcasting system that allows the AI agent to automatically discover which tools you have installed. Bio-babel-MCP thus serves knowledge through read-only tools that the agent calls; it neither plans nor executes.

We demonstrate the Bio-Babel-Toolkit chiefly on the R-to-Python package ecosystem. Specifically, we choose gpplot 2 ecosystem packages given its popularity and single cell trajectory, cell-cell communication analysis toolkits, such as Monocle 2 and NicheNet, earliest R packages developed for single cell biology. The rescontruction involves 5 layers (L0-L4) and is depth-first and rooted in grid, R’s abstract-layout substrate (Fig. 1c): rebuilding the grid first makes ggplot2 itself native, so new geoms, stats and scales can be written against Python internals, and higher-level methods built on the native grammar in turn. Above the L0 grid substrate sit L1 scales and gtable, L2 ggplot2, L3 grid-direct heatmap engines (pheatmap, ComplexHeatmap) and GOG extensions (ggrepel, ggalluvial, ggnewscale, patchwork, ggh4x), covering the custom Geom, Stat, operator, plot-composition and Facet mechanisms, and at L4, classical single-cell analytical tools such as Monocle 2/3^2,6^ and tradeSeq^7^, which then fully inherit the native visualization mechanisms in R. The result is a dependency-linked tree, each layer is resolved against the Python internals of the one below, and none calls back into R at runtime through a bridge such as rpy2. The reconstruction also re-roots each package on the target ecosystem’s native substrate rather than transliterating R’s, for example, the single-cell tools built on the community-standard AnnData^3^ container in place of Monocle 2’s bespoke R object container. The resulting stack therefore delivers not isolated packages carried from source to target language, but packages that form and enrich the target ecosystem.

We then measured fidelity against the R originals on three hierarchical tiers (Methods). Because the packages span diverse functions, we used multiple metrics, for example structural similarity (SSIM), Pearson correlation, overlap score and others (Supplementary Table 1). Overall, tier-resolved parity ranged from 62% to 100% across the reconstructed packages, with high pytest (72–98%) and documentation (64-95%) coverage (Fig. 2a). For ggplot2, Tier 1 verifies the numerical outputs of the data-transformation pipeline, passing 10/12 at max|Δ|≤1×10^−5^; the two misses are the kernel-density estimate (max|Δ|=1.3×10^−3^) and the intrinsically stochastic jitter position. Tier 2 (single Geoms) and Tier 3 (composed scenes) pass at 95% and 92%, respectively, with representative renders for ggplot2 and its underlying grid (Fig. 2b and Supplementary Fig. 3). For the bioinformatic tools, Monocle 2’s reconstructed components matched their R counterparts almost exactly (r = 0.999–1.000; Jaccard index = 0.961; adjusted Rand index = 1.00), and full-pipeline pseudotime agreed across the HSMM, lung, Paul and Olsson datasets (|ρ| = 0.98-1.00), the lowest being the Olsson state assignment (ARI = 0.78; Supplementary Fig. 4a). For tradeSeq, the substituted negative-binomial GAM backend reproduced the R fit (r = 0.826-0.999), as did predictSmooth fitted values and Wald tests (r = 0.908-0.995; Supplementary Fig. 4b).

**Figure 2.**
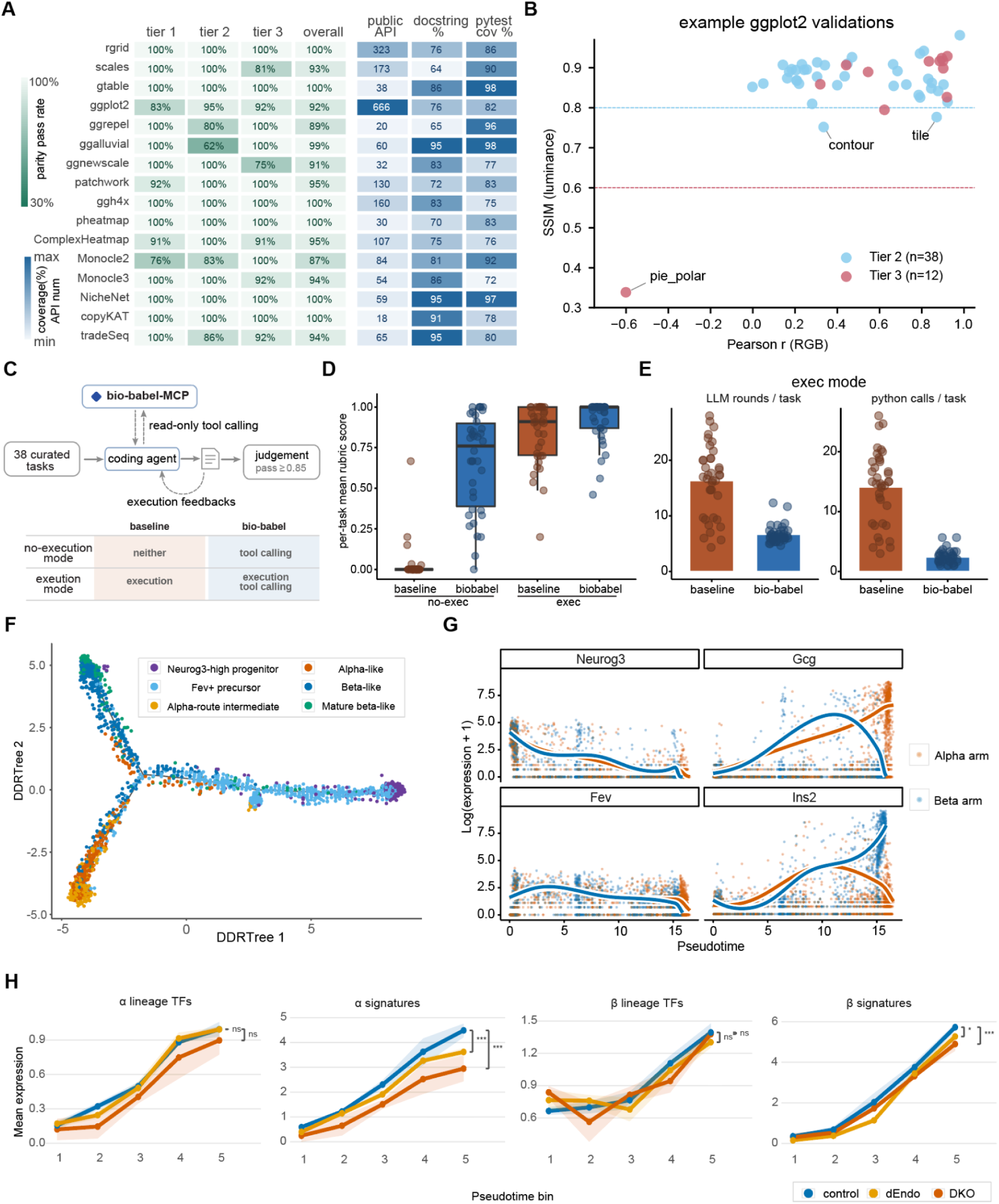
Validations and demonstrations of Bio-Babel framework. (**A**) Heatmaps of translation quality (including three tier validation scores and overall scores) and number of exported functions (public API) and coverage of docstrings and pytest for the re-implemented packages. (**B**) Scatterplot of the Pearson *r* v.s. SSIM score for tier 2 & tier 3 validations of ggplot2. Each point corresponds to one scene rendered by the original R and the new Python implementation, represented by structural similarity (SSIM, luminance) against pixel-wise Pearson correlation (RGB). Green and red dashed lines mark tier acceptance thresholds for the Tier 2 or Tier 3 validation, respectively; labelled points are the scenes that fail to pass the corresponding thresholds. (**C**) Schematic workflow for Bio-Babel-MCP benchmarking. Dashed lines indicate the two ablation settings. (**D**) Boxplot of the rubric scores on the Bio-babel-Bench: per-task mean rubric score for the two arms (baseline and Bio-Babel) under both evaluation modes (non-execution and execution). Boxes, interquartile range; centre line, median; whiskers, 1.5× IQR; points, per-task means. (**E**) Boxplot of LLM rounds or Python executions per task in execution mode. (**F**) DDRTree trajectory of 3,135 endocrine cells collected from nine embryos, coloured by manual cell annotation. Grey lines show the principal tree. The same colormap is used for panel G. (**G**) tradeSeq expression trends for Fev, Gcg, Ins2 and Neurog3 along alpha and beta branches. Points represent cells and lines show fitted trends. (**H**) The signature scores of pancreas α cells and β cells on their lineage transcriptional factors and marker genes along five even pseudotime bins.

Because these native packages follow Python idioms and depend on the Python ecosystem, they diverge from the originals by design, so an agent cannot fall back on prior knowledge of the R tools to call them correctly. To test whether the Bio-babel contract closes that gap, we built Bio-babel Bench, a new benchmark with 38 curated tasks that state a goal without naming the required API, graded by an independent model judge (Methods). We ran a coding agent in a 2×2 ablation experiment, crossing contract access with execution access (Fig. 2c). Without either, the agent almost never succeeded (failed 112 times out of 114 runs); the contract alone with no ability to execute code reduced this to 51%. When allowed to execute but given no contract, it still failed 36% of tasks; adding the contract cut failures to 16% (Fig. 2d) while shifting work from trial-and-error execution to context lookup, reducing both interaction rounds and code executions (Fig. 2e). This shift trades output tokens for input tokens spent on accurate lookups, at the cost of raised per-task cost (Supplementary Fig. 5).

Moreover, data analysis is increasingly done in dialogue with a coding agent: it writes and runs the code while the analyst steers the questions. To exercise both the building and interface sides of Bio-babel in that setting, we applied it to a published E15.5 mouse pancreas dataset spanning a graded loss of SWI/SNF catalytic activity^8^. Given the annotated data and statements of goals (Methods), the agent queried Bio-babel-MCP (Supplementary Fig. 6) and composed codes for trajectory analysis. monocle 2-python recovered the differentiation hierarchy: a Neurog3/Fev positive trunk bifurcating into Gcg^+^ alpha and Ins2^+^ beta branches (Fig. 2f, g and Supplementary Fig. 7a, b). As previously reported, neither knockout significantly altered the percentage of cells in alpha and beta clusters (Supplementary Fig. 7c). Interestingly, we found that alpha and beta signature scores rose along pseudotime but were attenuated with the severity of SWI/SNF loss, whereas the branch-defining transcription factors were retained (for example, Irx2 and Nkx6-1; Fig. 2h). This dissociation is consistent with the fact that the SWI/SNF complex is a permissive rather than an instructive factor. Kcnip4 was upregulated in both lineages, a shared response to SWI/SNF deficiency that may reflect remodelling of membrane excitability and stimulus–secretion coupling across both lineages under perturbations^9,10^ (Supplementary Fig. 7d).

Beyond the above demonstrated R-to-Python reconstruction, Bio-Babel is also extensible to other language pairs, requiring three components to be re-defined: the workflow steps, a set of language-specific skills and relevant scripts. For example, extending it to Python-to-C++ requires the addition of a sanitizer step that detects and removes undefined behaviour, scripts for C++ compilation and testing, and references covering C++ features and their common pitfalls (Supplementary Fig. 8a, b). We demonstrated the Python-to-C++ translation with the UMI-tools^11^. We then tested the C++ version of UMI-tools on the 10x Genomics PBMC 1K dataset, and recovered identical barcodes and identical UMI groupings (Supplementary Fig. 8c). Moreover, even without agentic optimization^12^, it consumed substantially less time or memory than the Python original across several commands, reflecting the intrinsic advantage of a compiled language (Supplementary Fig. 8d).

In summary, Bio-babel shows that a hierarchical ecosystem can be reconstructed faithfully in target languages, for example, from R to Python and that machine-readable operating knowledge is what further makes the reconstruction usable by agents. Pairing native, idiomatic implementations with the contract an agent needs to call them lets agents reliably extend, combine, and adapt existing pipelines, a prerequisite for the autonomous analysis workflows that computational biology is moving toward. The framework is not specific to graphics or to R to Python translation: it offers a route for tool developers to release native, multilingual versions of their methods, and for classic algorithms stranded in legacy languages to be carried into modern, widely used ecosystems while preserving the behaviour that made them worth keeping. Our reconstructed software ecosystem and Bio-Babel framework can be found here: https://github.com/Bio-Babel/. Future integration of Bio-Babel with PantheonOS (https://pantheonos.stanford.edu/), our agent-orchestration system, could further streamline translation and agent adoption across software packages and program languages.

## Methods

### Bio-Babel overview

Bio-Babel (<u>biobabel.stanford.edu</u>) is a framework for rebuilding scientific software ecosystem natively in another language and making it callable by agents, which contains four parts, namely Bio-Babel-toolkit, Bio-Babel-MCP, Bio-Babel-annotator, and Bio-Babel-Bench. Bio-Babel also ships a contract for each translated package to allow easy access by AI agents, such as Codex, Claude code, PantheonOS and others. Bio-Babel-Toolkit is the build side, driving coding agents to produce the translated packages, each a native member of the target ecosystem shipping a contract in its own source tree. Bio-Babel-annotator writes that contract, and Bio-Babel-MCP serves it to agents as a read-only surface. Here we migrate a hierarchical R ecosystem to Python, rooted in grid and ggplot2 as a native visualization foundation with classical single-cell tools built on top. We also demonstrated this framework extends to other language pairs, e.g. Python to C++, by re-specifying three components: the workflow steps, the reference skills and relevant scripts. Each part is detailed in the sections below.

### LLM agent-driven porting pipeline

Bio-Babel-toolkit is the LLM agent-driven porting system. Each port is produced by large language model (LLM)-based agents that follow a 13-step workflow. Throughout, the agents keep their state in a tree of report files and a machine-readable status YAML (YAML Ain’t Markup Language, a highly readable data formatting language especially useful for configuration files such as manifest files) file (*port_status.yaml*). This state is initialized at the outset, and it is re-read whenever an agent loses its context. The framework is extensible. Unless stated otherwise, however, porting in this manuscript refers to R-to-Python porting, and it proceeds through the following steps:

(1) Initialize the report tree, which contains the templates to sync the workflow state between local files and live agents;
(2) Establish the R and Python environments. Reuse those the user provides, which is the recommended practice, and otherwise create them;
(3) Audit the R package surface, including its exported symbols, S4/S3/R6 object systems, compiled (C/C++) sources, bundled and internal data, and test suite;
(4) Generate a feature checklist enumerating every export, each tracked as implemented, validated, or deferred;
(5) Classify and migrate the data assets;
(6) Decide the compiled-code strategy: reimplement in Python, wrap as a native extension, or skip when there are no compiled sources;
(7) Resolve the implementation of external dependency symbols. Reimplement, wrap an existing equivalent or analog packages, for example Python equivalent of R VGAM is the statsmodels packages;
(8) Build the translation and design map: normalize the API to Python conventions (snake_case for functions, methods, and properties; PascalCase for classes; R accessors as properties, replacement functions as explicit setters), select the target container (an established one such as AnnData where available, otherwise a purpose-built class), and lay out the modules;
(9) Parse the export surface into a dependency graph, partition it into dependency-ordered slices and implement the slices (see more in the next section);
(10) Validate the tutorials against R reference, skipped when the package ships no tutorials;
(11) Stage the remote data assets and generate the download registry, skipped when none are declared;
(12) Reproduce each R vignette as an executed tutorial notebook, skipped when there are no tutorials;
(13) finalize the documentation, including function/class signatures and tutorials if present.

### Graph-based optimization and parallel implementation

The code implementation of ported packages is guided by a LLM-based dependency analysis and graph partitioning (step 9). Firstly, the agent parses the package’s exports into a dependency graph, in which a directed edge from export A to export B means that the definition of A references B. This graph may contain cycles, so each strongly connected component (a set of symbols that define or refer to each other in a cycle or mutually recursive symbols) is contracted into a single node. The result is a directed acyclic graph. It is then partitioned into ordered slices, so that the unit tests of each slice depend only on slices implemented earlier. By default, the partition is computed by a cohesion-aware algorithm. This algorithm follows the dependency order and, at the same time, groups tightly coupled symbols together. It proceeds in three stages. First, it assigns each node a topological level, where level 0 holds the symbols that depend on nothing. This ordering fixes the slice order, and it guarantees by construction that every slice depends only on earlier slices. Second, it detects slices of densely connected symbols by weighted Louvain modularity maximization. It then groups the nodes into (level, slice) cells and greedily merges adjacent cells that share a slice, up to a size cap of 30 symbols. Finally, a cleanup pass merges any remaining small slices, splits any slice above the cap, and keeps every class together with its methods.

The method reasons only over exported functions and classes. Therefore, a manual slice that holds the data exporters, builders, and loaders is prepended. The slices are then implemented in order; independent slices may be optionally implemented concurrently by separate parallel sub-agents. In that case, each sub-agent’s edits are checked against the version-control diff for out-of-scope changes, and a slice is integrated only after it passes its unit tests and parity validation.

### Supervised orchestration and termination

A deterministic Python controller drives each port to completion, but it never writes code itself. Instead, it starts one coding-agent session per port and resumes that session each round, for up to 20 rounds. Each round begins by re-deriving the state from the local report files: the controller re-parses the reports and test results and converts them into a yaml file (*port_status.yaml*), and it sets the current phase to the first step that is not yet complete. It then runs diagnosis. If the port is unfinished, it resumes the agent and passes the remaining gaps. A port counts as complete only when four conditions hold together:

(1) every step is complete or skipped.
(2) the checklist is fully resolved, so that validated and deferred exports equal the total; a merely implemented export does not count.
(3) every tutorial is reproduced or validated.
(4) no diagnostic reports a gap.

Two safeguards keep this loop honest. The first is a stall detector. Between rounds, it compares a compact fingerprint of the state and the set of changed files. If the fingerprint is unchanged but files are still changing, the round counts as work in progress. If neither changes, a stall counter is incremented. The loop will then escalate to semantic diagnosis at two consecutive stalls and will abort for human intervention at three consecutive stalls. The second safeguard is diagnosis, which runs in three layers, namely structural, adversarial and semantic layers, and feeds *typed gaps* into the next prompt. The structural layer applies deterministic checks, such as whether every report is present, whether *pytest* passes. The adversarial layer runs once: it fans out multiple read-only sub-agents to trace each export’s call chains. The semantic layer runs on a stall, and it cross-references the local reports against the live code to flag claims that lack supporting evidence.

### Three-Tier validations for the re-implemented packages

We validated each reconstructed package against its R original. Tier 1 compares the numeric kernel of one function on a fixed input, such as when the graphic grid system converts a length between units. Tier 2 admits exactly one substituted layer, either a numerical backend, such as statsmodels for VGAM or scikit-learn for Rtsne, or the Python rendering engine drawing a single geom. Tier 3 runs a complete vignette end to end, so deviations from every layer compound, such as when monocle 2 performs the pseudotemporal ordering of single cells from a dataset.

Reference values were generated under each original’s pinned R environment and frozen, so no comparison invokes R. Where a step is stochastic, we load R’s own draw rather than re-sampling. An observable is one scalar comparison against a named R reference. Per-element comparisons are counted individually. A probe over a vector or matrix counts as one comparison, using its worst-case deviation, while a rendered figure counts as one comparison, using the structural similarity (SSIM) between the two images computed on luminance. In total, this gives 1,510 scored observables over the 16 packages (DDRTree being validated within monocle 2, see Code Availability for the full package list). We defined seven metric families (Supplementary Table 2) spanning the quantities these packages produce and assigned every observable to one of them. We divide the seven metrics into two groups. For similarity metrics (correlation, clustering, set overlap, and image similarity), thresholds become more permissive across tiers because higher tiers allow greater divergence. For exact-reproduction metrics (numeric deviation, resolved colour, and structural equality), the threshold stays the same across all tiers because these quantities must be reproduced accurately at every tier.

### Bio-babel-MCP protocol design

Each package carries a contract (rules and instructions) directory (*_biobabel*/) inside its source tree to facilitate interactions with agents. This directory declares, in YAML, everything an agent needs to call the package correctly, and a factory assembles its files into a single manifest, a configuration file that acts as a blueprint. The manifest holds six kinds of queryable objects:

(1) A symbol records a callable’s signature and parameters, the state it requires and writes;
(2) A concept states a package invariant, and in the context of R to Python translation, it gives a dual mental model, phrased once for an R user and once for a Python user;
(3) An idiom pairs a recommended pattern with a runnable code template;
(4) An anti-pattern instead pairs a discouraged pattern with the abstract-syntax-tree (AST) detector that recognizes it and with the idiom that replaces it;
(5) A workflow is a linear plan of (symbol, purpose) steps;
(6) and a template is an end-to-end script skeleton.

To build a bridge between AI agents and the packages Bio-Babel converts, we developed a Bio-Babel Model Context Protocol (MCP) server, namely, Bio-Babel-MCP, which defines the rules that allows agents to read and understand the “contract”. Instead of relying on a central registry, e.g., app store, discovery of the package is handled locally. The server enumerates a specific Python entry-point group, allowing each installed port to advertise its manifest. Consequently, the agent only sees the packages the user has explicitly installed, and any package lacking a defined contract remains completely invisible.

### Automated contract generation

To automate contract generation, we developed an LLM annotator agent (Bio-Babel-annotator), Bio-Babel-annotator, to generate a contract for each package. The contract class of a package strictly defines the agent’s expected output. While every class requires symbol contracts detailing its complete public surface, specific package types carry additional requirements: analysis packages must specify workflows, whereas grammar packages must outline concepts, idioms, and anti-patterns. This contract class is either provided by the operator or inferred autonomously by the agent via the public API.

Starting from an empty *_biobabel*/ directory, a controller builds the contract. It initializes one agent session, and resumes it each round with the gaps the previous round left. The loop stops when the contract validates clean, when a round changes nothing, or at a round cap. Validation is deterministic and checks seven conditions:

(1) the contract satisfies the manifest schema;
(2) every public export carries a symbol contract, so that coverage will not silently shrink;
(3) all cross-references resolve;
(4) every documented parameter exists in the implementation’s real signature;
(5) no workflow step requires state that only a later step produces;
(6) each anti-pattern’s detector fires on its own negative example and stays silent on the positive one;
(7) the server discovers the installed package’s manifest.

If we cannot decide the findings mechanically, it will stay advisory and never drive repair. These checks establish the contract’s form, coverage, and consistency; the agent itself writes the interpretive content from the package’s source code, tests, and documentation.

### Benchmark the Bio-babel-MCP for agent coding

To benchmark the agent coding capability of Bio-babel-MCP contract layer, we firstly designed Bio-Babel-Bench (https://github.com/Bio-Babel/Bio-Babel-Bench), which contains 38 intent-level coding (the process of instructing AI with a high-level goal, leaving agent to autonomously generate the exact implementation) tasks. Each task states a goal without naming the required API and specifies the expected artifacts, difficulty level, and a 100-point grading rubric. The tasks cover single API elements, common API compositions, and realistic multi-step workflows. Because a vision language model doesn’t provide mathematical precision and non-determinism, in this Bench, the grader doesn’t inspect rendered images, visualization tasks must produce machine-readable artifacts containing the resolved values.

We used GPT-5.4 as the coding agent in a controlled (2*2) ablation with two factors: contract access and execution access, where the first factor has two arms (baseline arm and treatment arm) and the second factor has two regimes (non-execution regime and execution regime). The baseline arm received only workspace tools, whereas the treatment arm received the same tools plus access to the Bio-Babel-MCP contract server. Under the no-execution regime, neither arm had an execution tool; the agent produced one solution file, which the harness executed once before grading. Under the execution regime, both arms could run and revise their code for up to 40 tool-use rounds. The former measures one-shot API recall, whereas the latter measures the efficiency of reaching a correct solution through iteration. All other settings were identical across conditions. Contract leakage was prevented structurally. Both arms executed code in an environment built from a source tree, the complete directory structure containing all the source code files for a project, containing no contract files. The contract server ran in a separate environment built from a parallel source tree containing the contracts and was launched only in the treatment arm.

To avoid self-judgement, we used GPT-5.5 as an independent judge. It received the agent’s final response, source code, and generated text, JSON, or CSV artifacts. Rubric points were summed and normalized to ([0,1]), with scores of at least 0.85 counted as passes. For each run, we recorded the score, pass outcome, execution and contract-lookup calls, token usage, interaction rounds, elapsed time, and cache-aware cost at the subject model’s list price. Contract-lookup tokens were included in the cost. Each task-by-arm-by-regime condition was repeated three times. Results were aggregated by arm and regime, and we report score, pass rate, tool usage, cost, treatment-minus-baseline differences, and pass rates by library (Supplementary Table 3).

### Computational analyses of pancreatic lineages under SWI/SNF subunit losses

We chose a high quality single cell dataset to demonstrate Bio-Babel’s capability in enabling data analyses with the reconstructed software ecosystem that leads to novel biological discoveries. Processed single cell RNA-seq data from GSE248369 is downloaded from GEO and is converted into an AnnData object (h5ad format for storage). The data contains 5,196 manually annotated E15.5 pancreatic cells and 23,308 genes from ten embryos: three control, four dEndo (*Brg*^Δ*endo*^ ; *Brm*^+/−^ ) and three DKO (*Brg*^Δ*endo*^ ; *Brm*^−/−^ ) embryos.

These genotypes represent progressive loss of the BRG1 and BRM ATPase subunits of the SWI/SNF chromatin-remodelling complex in the pancreatic endocrine lineage.

In this analysis, we provide the coding agent (Claude code) with the h5ad file and a prompt describing the analytical goals, available at https://github.com/Bio-Babel/Bio-Babel-pancreas. The coding agent queried Bio-babel-MCP 19 times for workflow and API guidance and successfully reconstructed the alpha/beta cell trajectories automatically. It then independently generated, executed an analysis using monocle 2-python with DDRTree-python for trajectory inference and BEAM for lineage-specific gene expression testing, tradeSeq-python for pseudotime-dependent expression modelling and ggplot2-python for visualization. Finally, it gave examined outputs and a preliminary report with interpretations and recommendations for follow-up analysis.

We reviewed the resulting notebook and preliminary report, and performed targeted follow-up analyses. Monocle 2-python inferred the trajectory with DDRTree from normalized expression of 291 cell-type annotation-guided ordering genes, comprising the top 60 one-versus-rest genes for each of six retained annotations plus curated endocrine markers after filtering low-detection and technical genes. All 422 cells from DKO2, one of the three DKO embryos, were excluded because its transcriptome mapping rate was 46.7%, compared with 64.5–80.7% for the other libraries. The final trajectory contained 3,135 progenitor, precursor, α-like and β-like cells from nine embryos. BEAM tested 16,888 genes detected in at least 20 cells using negative-binomial models. The full model was

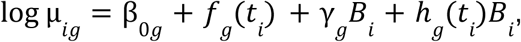

and the reduced model was

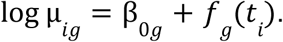

Here, *i* indexes cells and *g* indexes genes; µ*_ig_* is the expected size-factor-normalized expression of gene *g* in cell *i*; β_0*g*_ is the gene-specific intercept; *ti* is pseudotime for cell *i*; *B_i_* is an indicator of branch assignment for cell *i*; *f_g_* (*t_i_* ) is the common pseudotime trend; γ*_g_* is the branch main effect; and ℎ*_g_* (*t_i_* ) is the branch-specific deviation from the common trend. γ*_g_B_i_* indicates the constant branch offset for gene *g* while ℎ*_g_* (*t_i_* )*B_i_* a branch-by-pseudotime interaction. The full and reduced models were compared using a likelihood-ratio test. Lineage-associated transcription factors (Arx, Irx1, Irx2 and Pou3f4 for α cells; Pdx1, Nkx6-1 and Mafa for β cells) and terminal effectors (Gcg and Ttr for α cells; Ins1, Ins2 and Iapp for β cells) compared between genotypes were selected from 100 highest-ranked BEAM genes. The 1,000 highest-ranked BEAM genes were combined with the 400 most variable genes and a curated marker panel, yielding 1,116 unique genes for tradeSeq analysis. Claude initially fitted genotype-aware tradeSeq models for 150 of these genes. We extended the same analysis to all 1,116 genes using condition-aware models with inferred sex as a covariate. Raw counts were fitted using six-knot negative-binomial generalized additive models:

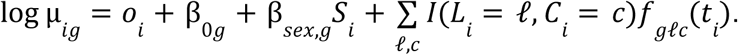

Here, µ*_ig_* is the expected raw UMI count of gene *g* in cell *i*; *o_i_* is the cell-specific trimmed mean of M-values or TMM log-offset; β*_og_* is the gene-specific intercept; *S_i_* is the inferred embryo sex indicator for the embryo containing cell *i*, and β*_sex_*_, *g*_ is its gene-specific coefficient; *l* and c index lineage and genotype; *L_i_* and *C_i_* denote the lineage assignment and genotype of cell *i*, *I*(·) is the indicator function; and *f_glc_* (*t_i_* ) is the gene-, lineage- and genotype-specific pseudotime smoother. Genotype effects were tested using lineage-specific pairwise Wald contrasts of the fitted smoother coefficients. Genes were retained if their control mean log1p(CP10K) was at least 0.1 in both trajectories. For each mutant–control contrast, a concordant cross-lineage candidate required the same sign of mean fitted log2FC across 100 pseudotime points and a Benjamini–Hochberg-adjusted q < 0.05 for the |log2FC| > 0.5 effect-threshold test in both α and β trajectories. Candidates shared by both mutants passed these criteria in both contrasts. Benjamini–Hochberg correction was applied separately within each lineage–contrast test. These cell-level tests were used for candidate ranking rather than biological-replicate-level inference.

### Extensibility of Bio-Babel-toolkit to other language pairs

Extending the framework to a new language pair requires re-specifying three language-specific components: the workflow steps, the checks and the skills. The steps keep the spine and add what the target language requires. Python-to-C++ adds a sanitizer step. Undefined behaviour in C++ compiles and runs while producing plausible output, so an uninitialised read can by itself produce a matching comparison. The C++ version is therefore required to be free of undefined behaviour before any parity result is recorded. The checks are what the controller reads to decide whether a step is finished. Configuring, compiling and testing can each fail independently, so they are checked separately. A successful configuration is also the evidence that the mapped C++ dependencies link. The skills hold the target-language references and scripts. The references cover Python-to-C++ translation patterns, build and dependency handling, undefined behaviour, and parity. The scripts verify the toolchain, generate the project and record state.

The C++ version was validated on the same three tiers, against the running Python original, using the Python package’s own test fixtures, adversarial inputs and real data. For UMI-tools, 121 exports and CLI subcommands were validated with none deferred, and 216 comparisons passed. Efficiency was then measured on the 10x Genomics PBMC 1K dataset, comprising 66,601,887 read pairs. The whitelist, extract, group, dedup and count subcommands each returned byte-identical output under both versions. Timings were single-threaded, with one discarded. Warm-up followed by three repeats in which the two versions ran back to back on the same node.

## Data Availability

The single-cell RNA-seq data re-analysed here are publicly available from the Gene Expression Omnibus under accession GSE248369; the dataset used to validate and benchmark the C++ version of UMI-tools is the 10x Genomics 1k PBMCs from a Healthy Donor (v3 chemistry) dataset, released with Cell Ranger 3.0.0 (10x Genomics, 2018) and available at https://www.10xgenomics.com/datasets/1-k-pbm-cs-from-a-healthy-donor-v-3-chemistry-3-standard-3-0-0; all other datasets used for validation are distributed with the corresponding R packages or their original publications.

## Code availability

The Bio-babel project website is available at <u>biobabel.stanford.edu</u>. Core Bio-Babel packages include: Bio-Babel-Toolkit (https://github.com/Bio-Babel/Bio-Babel-Toolkit), Bio-Babel-MCP (https://github.com/Bio-Babel/Bio-Babel), Bio-babel-bench for MCP ablation benchmark experiments (https://github.com/Bio-Babel/Bio-Babel-Bench), Bio-Babel-annotator (https://github.com/Bio-Babel/Bio-Babel-annotator). And, the scripts and other necessary materials to reproduce the analysis in Figure 2 (https://github.com/Bio-Babel/Bio-Babel-pancreas). Bio-Babel-Toolkit and Bio-Babel-annotator will be public upon publication.

Besides, we also released all packages produced with the Bio-babel muti-agents framework:

(1) rgrid-python (https://github.com/Bio-Babel/rgrid-python);
(2) scales-python (https://github.com/Bio-Babel/scales-python);
(3) gtable-python (https://github.com/Bio-Babel/gtable-python)
(4) ggplot2-python (https://github.com/Bio-Babel/ggplot2-python);
(5) patchwork-python (https://github.com/Bio-Babel/patchwork-python);
(6) ggrepel-python (https://github.com/Bio-Babel/ggrepel-python);
(7) ggalluvial-python (https://github.com/Bio-Babel/ggalluvial-python);
(8) ggnewscale-python (https://github.com/Bio-Babel/ggnewscale-python);
(9) ggh4x-python (https://github.com/Bio-Babel/ggh4x-python);
(10) pheatmap (https://github.com/Bio-Babel/pheatmap-python);
(11) ComplexHeatmap (https://github.com/Bio-Babel/ComplexHeatmap-python);
(12) Nichenet-python (https://github.com/Bio-Babel/Nichenet-python);
(13) Copykat-python (https://github.com/Bio-Babel/copykat-python);
(14) DDRtree (https://github.com/Bio-Babel/DDRTree-python);
(15) monocle 2-python (https://github.com/Bio-Babel/monocle 2-python);
(16) Monocle 3-python (https://github.com/Bio-Babel/Monocle3-python);
(17) tradeSeq-python (https://github.com/Bio-Babel/tradeSeq-python);
(18) Python2C++ package, UMI-tools-cpp (https://github.com/Bio-Babel/UMI-tools-cpp);

## Acknowledgements

This work is supported by Laude Moonshot Seed Grant, the Pantas And Ting Sutardja Foundation, the Wu Tsai Neurosciences Institute Big Ideas in Neuroscience Program, MorPhiC consortium (U01 HG013176), R00 or NIH Pathway to Independence Award (5R00HG012887-04), DP2 or NIH Director’s New Innovator Award (DP2HG014282-01).

