## supplementary_file for "Bio-Babel: autonomous cross-language reconstruction of agent-ready computational biology ecosystems"

### Supplementary Figures

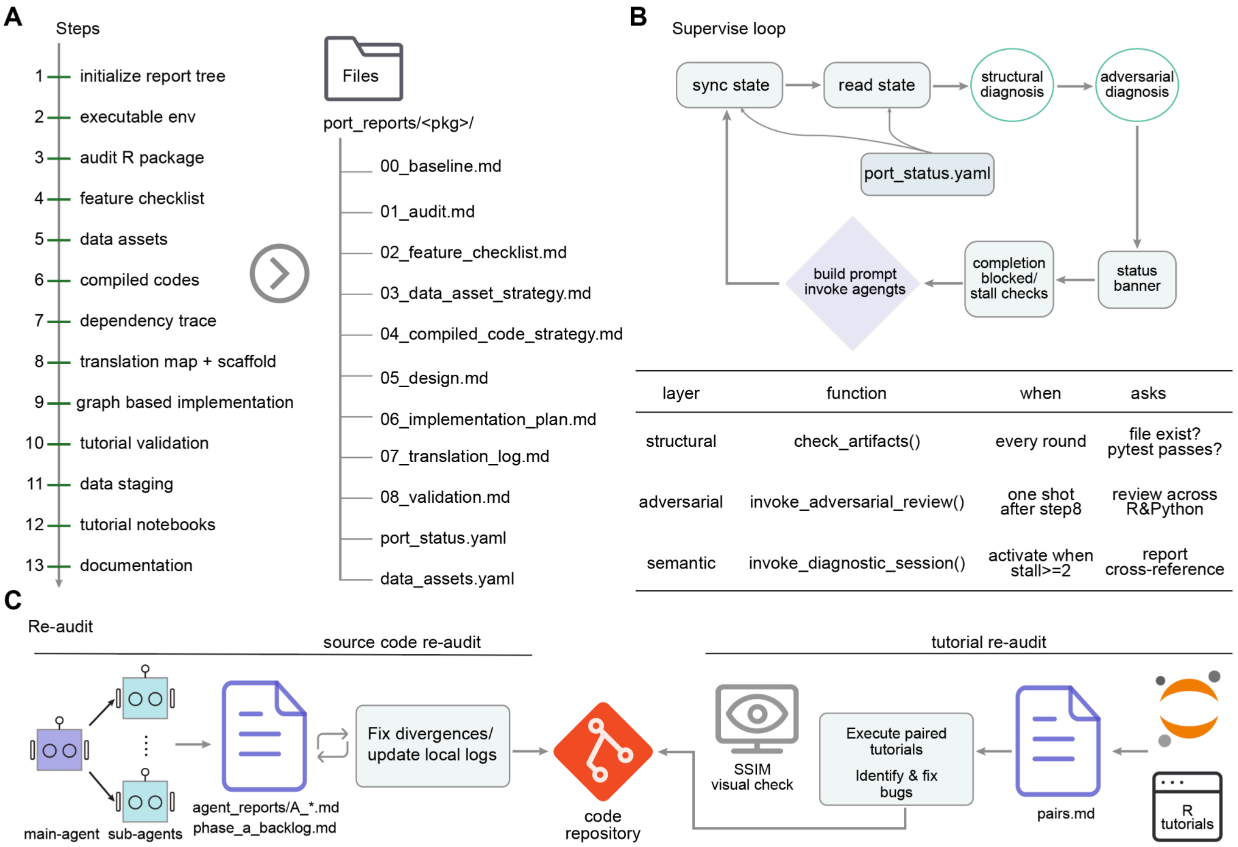


**Supplementary Figure 1** | The details behind Fig. 1b. (**A**) The detailed 13-step Bio-Babel-Toolkit workflow (left) and the durable report file tree structure within the port_reports/<pkg>/ folder which every step writes into (right). (**B**) A supervised loop controls the 13-step workflow, which on the one hand syncs the state between live agents and local reports, on the other hand advances the workflow based on the state. (**C**) Iterative re-audit process. It includes two parts: source code re-audit and tutorial re-audit. In the source-code re-audit, a main agent fans out parallel read-only sub-agents that audit each exports against its original, writing findings to agent_reports/A_*.md; in the tutorial re-audit, tutorials are executed and compared. Together, based on the audit, the agents further fix residual bugs iteratively.

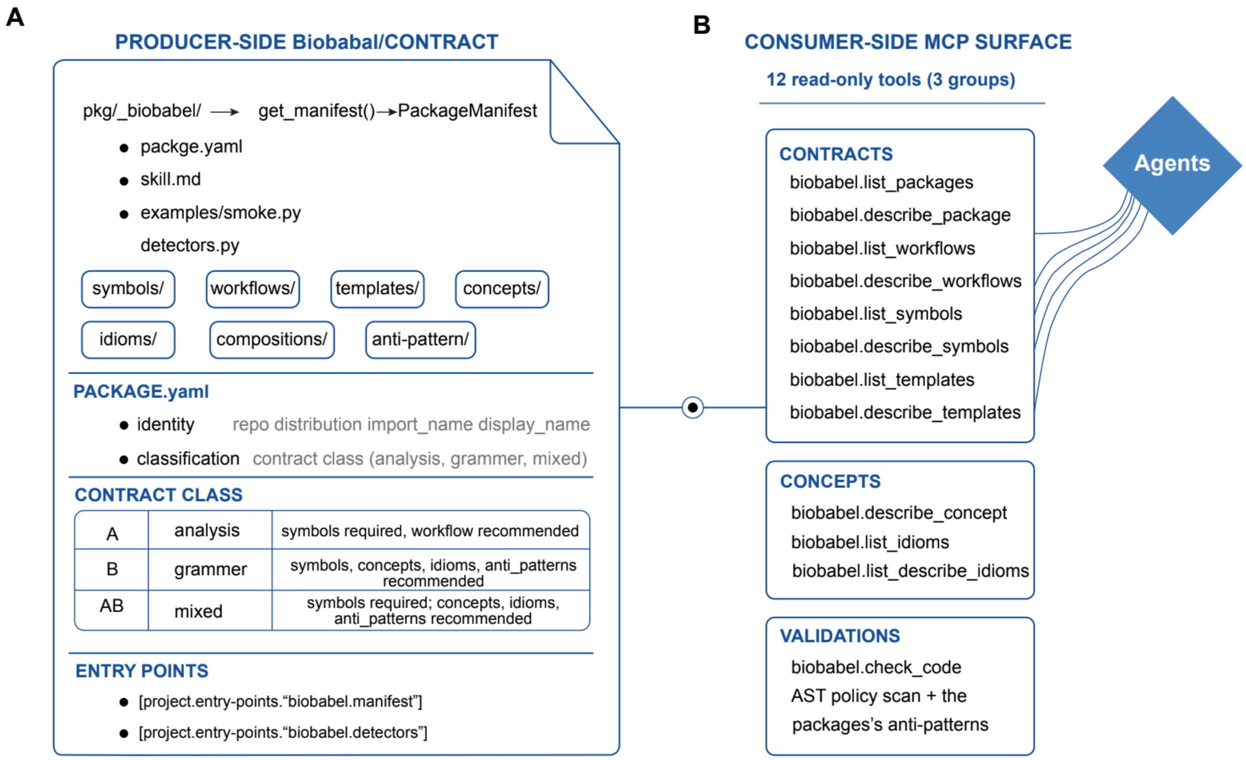


**Supplementary Figure 2** | The internal contract and the external MCP surface (expands Fig. 1d). (**A**) The internal _biobabel/ contract shipped inside every package (**B**) The MCP surface to external agents, e.g. Claude Code, Codex, PantheonOS, etc., 12 read-only MCP tools in three groups. The agent consumes these tools; Bio-Babel-MCP neither plans nor executes.

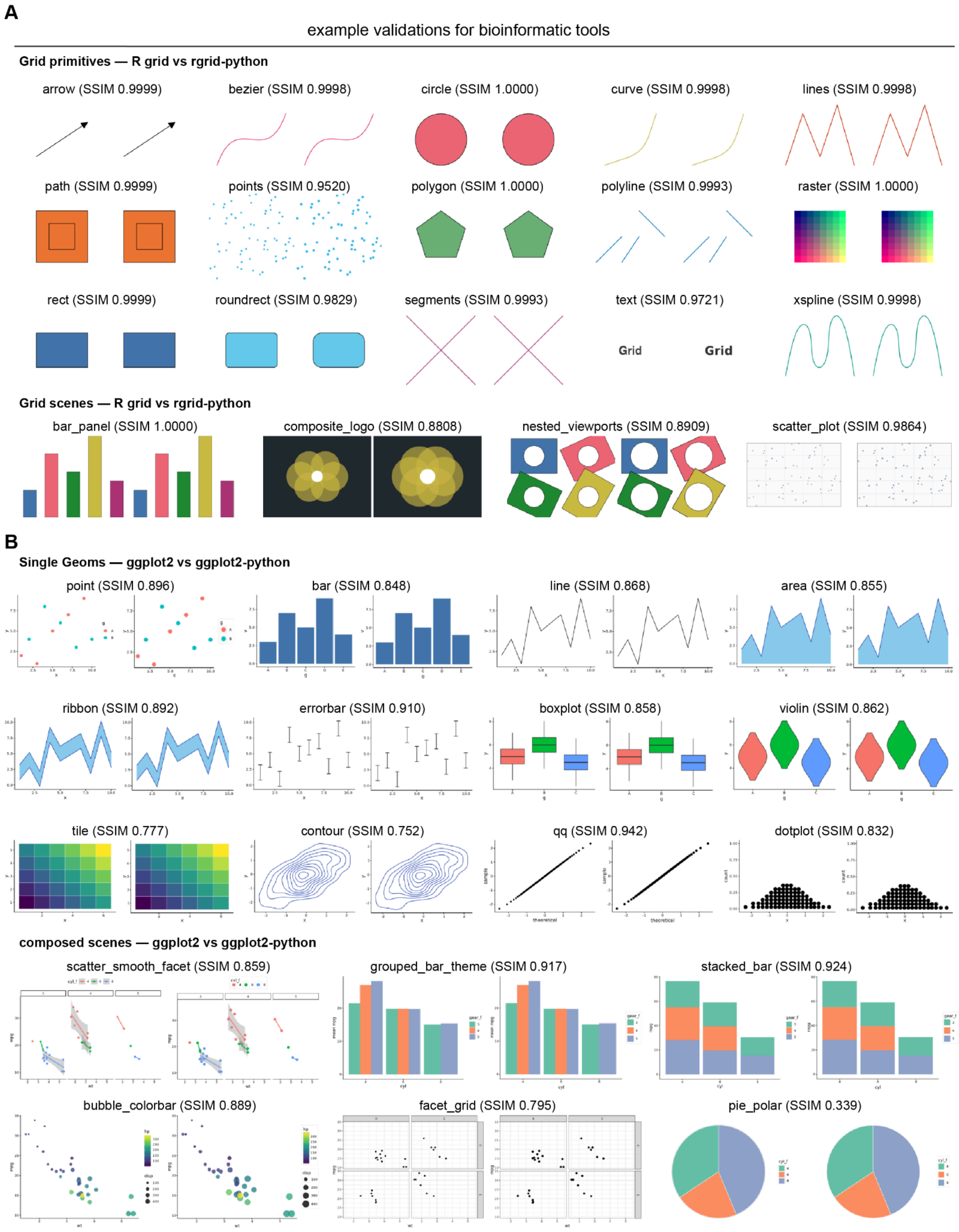


**Supplementary Figure 3** | Visual parity examples between R packages and translated Python counterparts for the package of graphics substrate (grid) and that of the grammar of graphics built on it (ggplot2). (A) R grid versus rgrid-python: 15 drawing primitives and 4 composed scenes. Representative renders are shown side by side for each package and every pair is quantified by the structural similarity index (SSIM, luminance channel). Same as for panel B. (B) R ggplot2 versus ggplot2-python: 12 single geometries and 6 composed scenes. The lowest-scoring pairs (tile and pie_polar) reflect legitimate rendering differences (anti-aliasing of dense marks; the polar-coordinate pie_polar scene), not content errors.

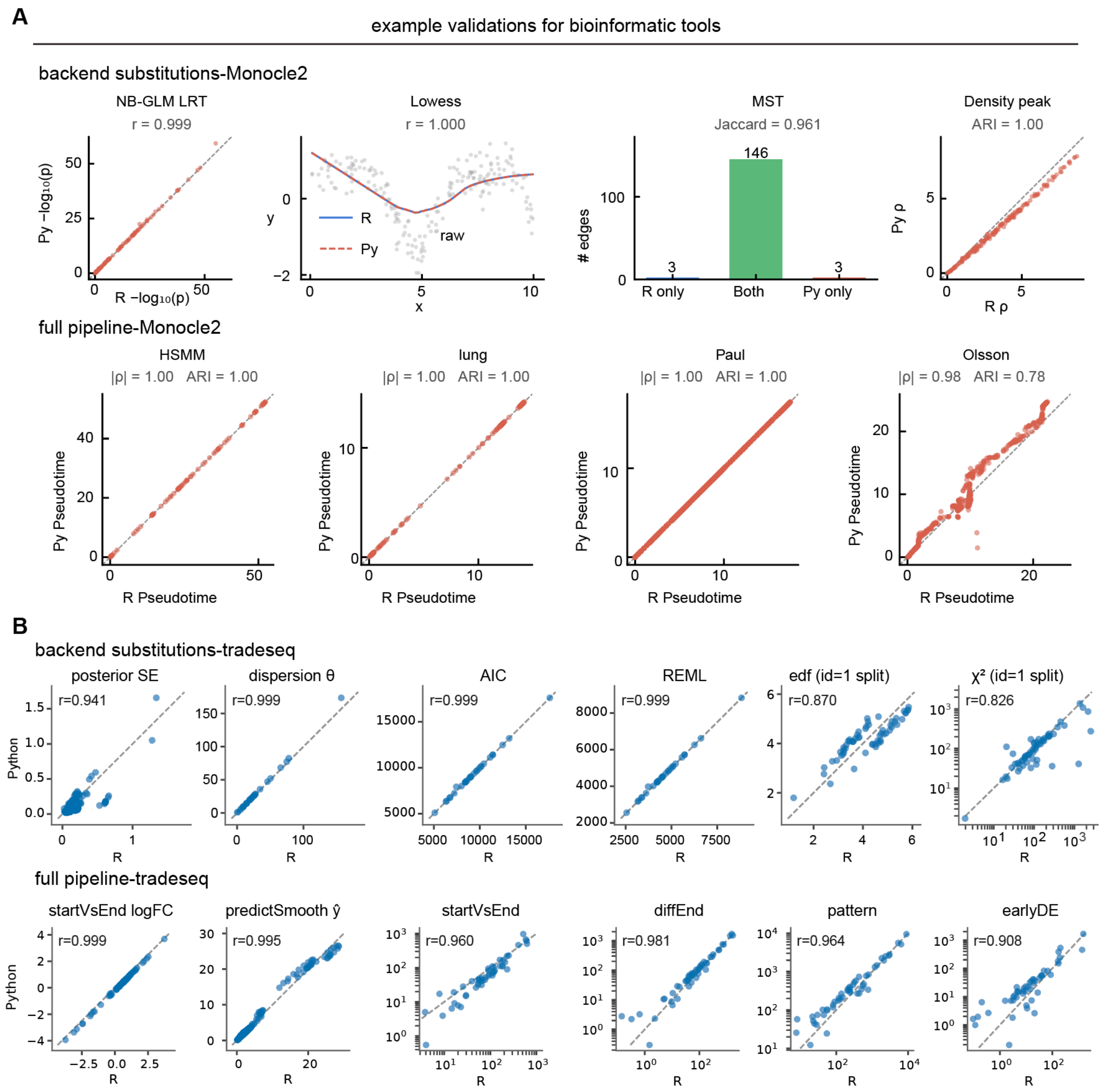


**Supplementary Figure 4 | Visual parity examples of bioinformation tools (monocle 2 and the tradeSeq) translation accuracy**. (**A**) Monocle 2 translation visual parity validation. Algorithms with substituted backends and full-pipeline results were compared between python packages and original packages. Same as for panel B. Top: comparisons of algorithms with backends substituted — the negative-binomial generalized linear model or GLM likelihood-ratio test, the lowess smoother, the minimum spanning tree and density-peak clustering (statsmodels vs R VGAM); Bottom: the scatterplot of full pseudotime analyses pipeline comparison shows agreements across four datasets — HSMM, lung, Paul and Olsson. (**B**) Same as in panel **A** but for tradeSeq translation visual parity validation. Top: the scatterplot of various statistics of tradeSeq under different backends, the negative-binomial generalized additive model or

GAM backend (statsmodels NB-GLM vs R mgcv), compared per gene : posterior standard errors, dispersion θ, AIC, REML score, the smoother's effective degrees of freedom (edf) and $\chi^2$ statistic (top). Pearson correlation r is listed for the comparison, same as for the bottom row. Bottom: the scatter plot of the downstream fitted start-to-end log-fold-change (startVsEnd logFC) and smoother curve (predictSmooth $\hat{y}$), and the Wald statistics of differential-expression tests (startVsEndTest, diffEndTest, patternTest, earlyDETest; log–log), across genes between R (x-axis) and translated Python implemented (y-axis)

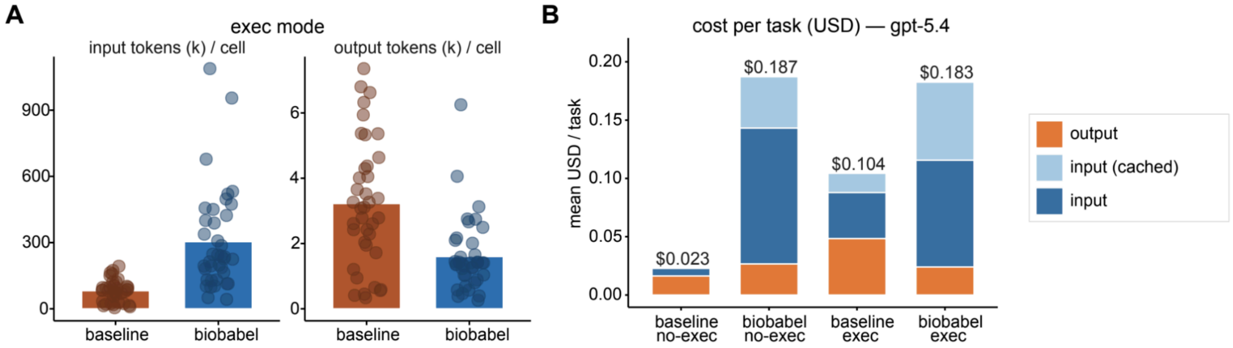


**Supplementary Figure 5** | Token and dollar cost of the ablation experiment. (**A**) mean token usage (input tokens and output tokens) per task for both arms in execution-mode. Each point represents a task,repeating 3 times (**B**) Mean US-dollar cost per task (gpt-5.4 pricing) across the four mode-by-arm cells, decomposed into output, cached-input and input tokens.

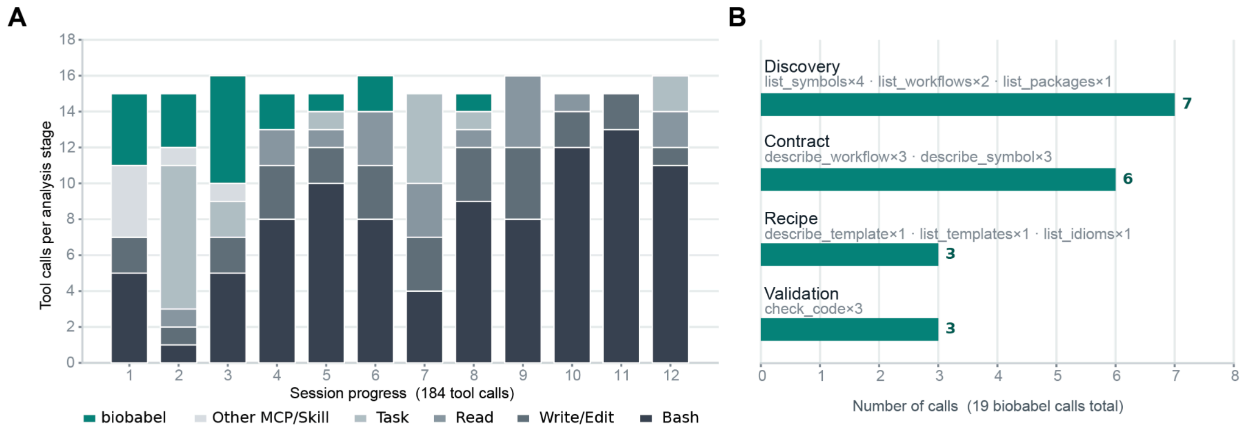


**Supplementary Figure 6 | Analysis of Bio-Babel-MCP invocation in an analysis example**. Tool-call profile of one end-to-end session that used the Bio-Babel-MCP when using agent to analyze the GSE248369 dataset. (A) The 184 tool calls across 12 sections in the analysis notebook, colored by tool class, with Bio-Babel (teal) stacked at the top of each bar. (B) The 19 Bio-Babel calls grouped into four functional roles (Discovery, Contract, Recipe, Validation); bar length gives the number of calls, with the constituent MCP tools listed above each bar.

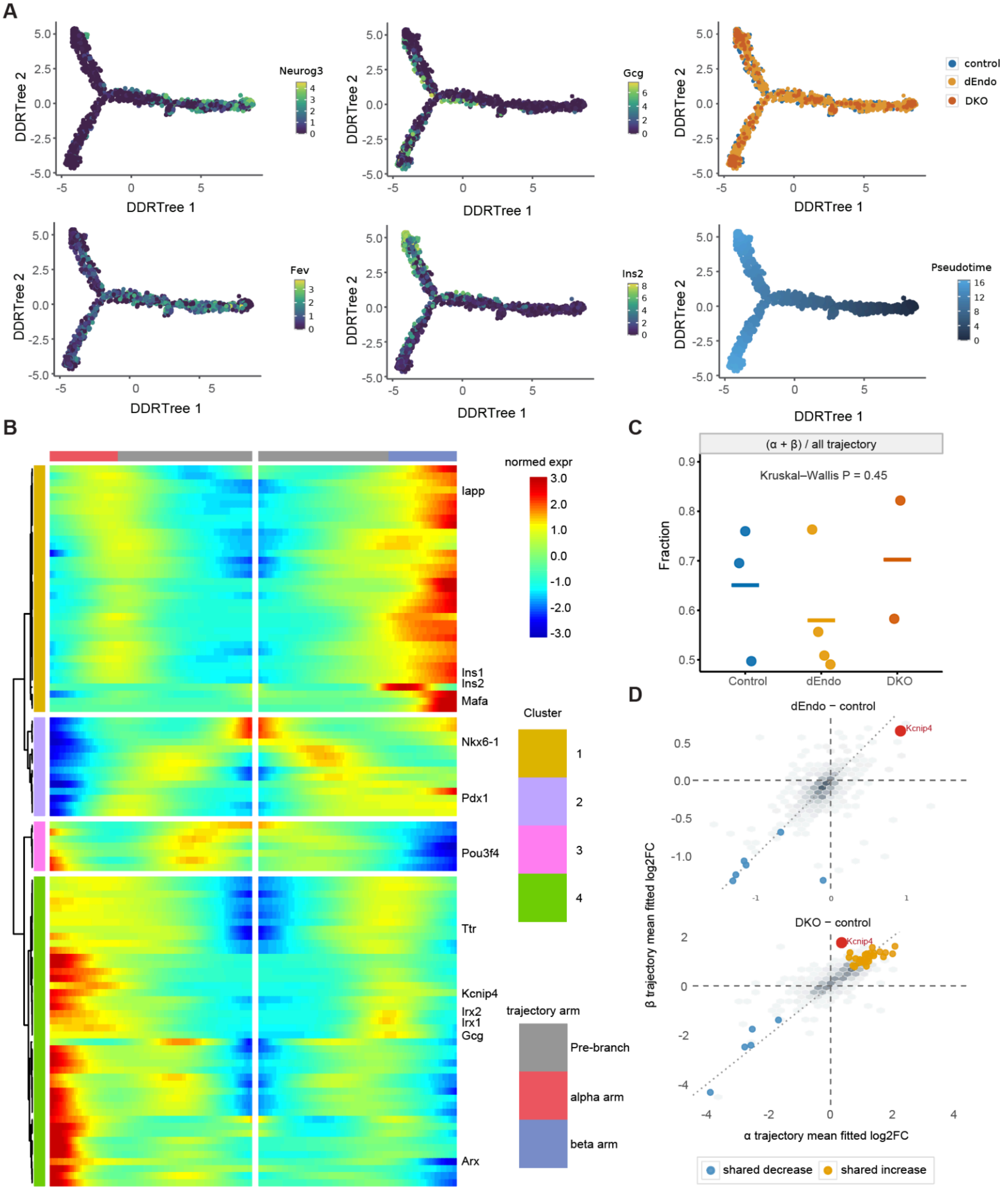


**Supplementary Figure 7** | trajectory analysis with monocle 2-python and tradeSeq-python. (**A**) DDRTree projections coloured by Neurog3, Fev, Gcg or Ins2 expression, genotype, or pseudotime. dEndo: *Brg1*$^{\Delta endo}$;*Brm*$^{+/-}$, DKO: *Brg1*$^{\Delta endo}$;*Brm*$^{\Delta endo}$. (**B**) BEAM heatmap of top 100 genes differing between the alpha and beta branches (q<0.05). Colours show row-scaled expression; annotations indicate gene programs and clusters. (**C**) Fraction of cells fallen along the α/β branch for individual embryos (control, n=3; dEndo, n=4; DKO, n=2). Horizontal lines show group means; no significant difference was detected (Kruskal–Wallis P=0.425). (**D**) Comparison of differential gene expression along the α- and β-cell trajectories in dEndo (top) and DKO (bottom) relative to their controls.

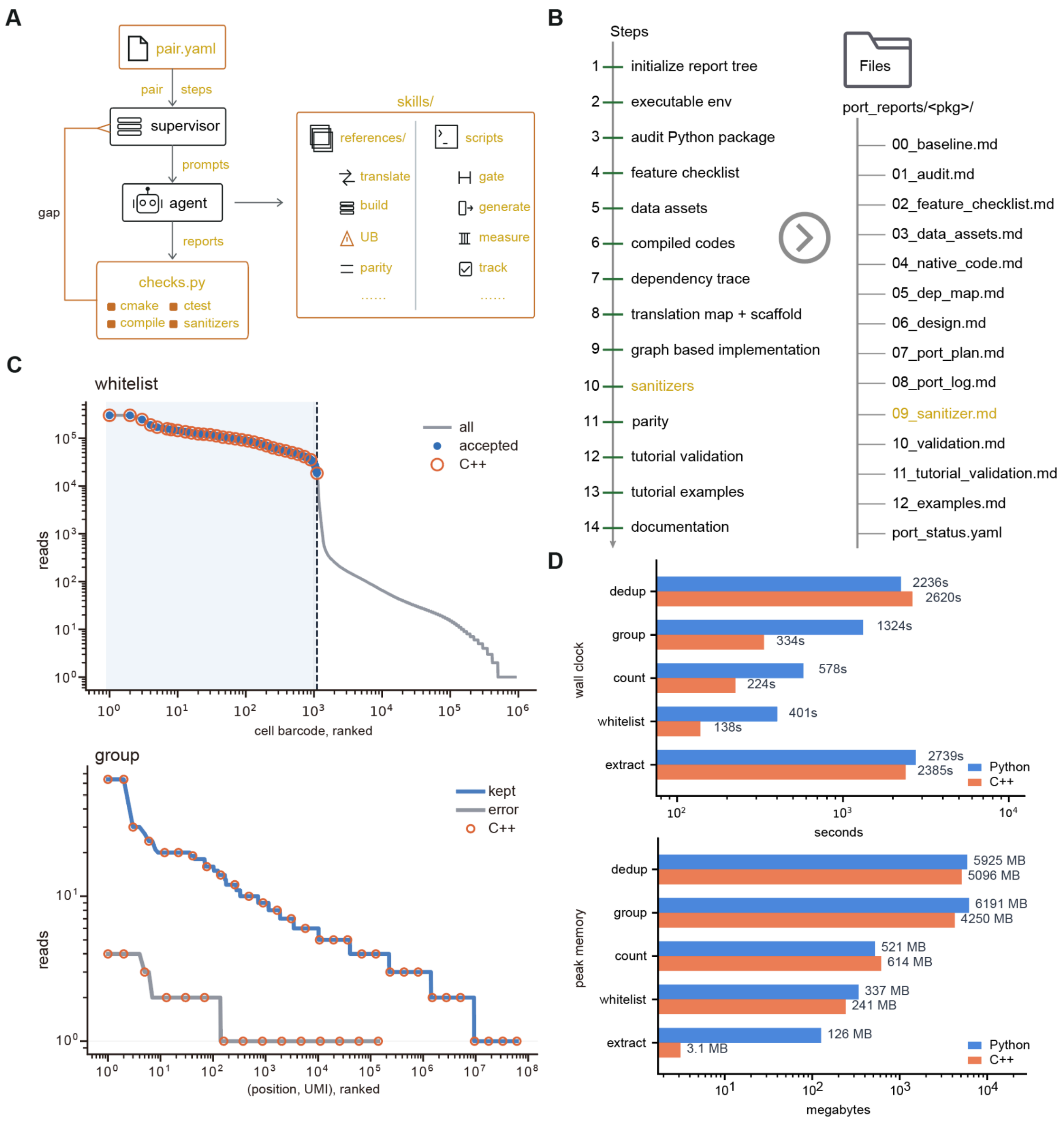


**Supplementary Figure 8** | Demonstration of the extensibility of Bio-Babel-Toolkit on a new language pair. (**A**) The schematic workflow to extend Bio-Babel-Toolkit to a new language pair, taking python2C++ as an example: (1) pair.yaml declares the steps of workflow, which could be adjusted accordingly based on different language features. In this case, we add a sanitizer step that specifically cleans the undefined behavior (UB) in C++; (2) checks.py decides when a step is done, returning the gaps that drive the next round, which for C++ are cmake configure, compilation, ctest and the sanitizer build; and (3) the skills that hold both the references for the new language and the scripts the agent runs. The references introduce the target language to the agent, covering for example how Python constructs and NumPy arrays map onto C++ types and containers, how a cMake project resolves and links its dependencies, which forms of undefined behaviour C++ admits and which sanitizer builds expose them, and how a differential comparison against the Python original is recorded and reproduced. The scripts are what the agent runs at each step, for example verifying that the C++ toolchain and the Python reference environment are usable, generating the CMake project, running the differential comparisons, and recording progress on disk. (**B**) The steps in python2C++ workflow and its corresponding report tree. Orange text marks the step, and its report, that are unique when compared with the R2python workflow. This is the sanitizer step, which proves the C++ version is free of undefined behaviour before any parity result is recorded. (**C**) Result comparisons between the C++ version and original Python version on UMI-tools functions with 10x pbmc_1k_v3 dataset. The whitelist command separates real cells from empty droplets; the group command decides which reads came from the same molecule with UMI corrections. Both of the results on the two commands are identical. (**D**) Runtime (wall-clock time) and peak memory use (maximum resident set size, RSS) for the C++ and Python versions of UMI-tools. Values are medians of three interleaved, single-threaded runs; black ticks show the minimum–maximum range when it exceeds 2% of the median.
